# A new method for detecting mixed *Mycobacterium tuberculosis* infection and reconstructing constituent strains provides insights into transmission

**DOI:** 10.1101/2024.04.26.591283

**Authors:** Benjamin Sobkowiak, Patrick Cudahy, Melanie H. Chitwood, Taane G. Clark, Caroline Colijn, Louis Grandjean, Katharine S. Walter, Valeriu Crudu, Ted Cohen

**Affiliations:** Department of Epidemiology of Microbial Disease, Yale School of Public Health, 60 College Street, New Haven, CT, USA; Department of Infection, Immunity and Inflammation, Institute of Child Health, University College London, London, UK; Division of Infectious Diseases, Department of Internal Medicine, Yale School of Medicine, New Haven, CT, USA; Faculty of Infectious and Tropical Diseases, London School of Hygiene and Tropical Medicine, UK; Faculty of Epidemiology and Public Health, London School of Hygiene and Tropical Medicine, UK; Department of Mathematics, Simon Fraser University, 8888 University Drive West, Burnaby, BC, Canada; Division of Epidemiology, University of Utah, Salt Lake City, UT, USA; Phthisiopneumology Institute, Strada Constantin Vârnav 13, Chisinau, Republic of Moldova

## Abstract

**Background:** Mixed infection with multiple strains of the same pathogen in a single host can present clinical and analytical challenges. Whole genome sequence (WGS) data can identify signals of multiple strains in samples, though the precision of previous methods can be improved. Here, we present **MixInfect2,** a new tool to accurately detect mixed samples from *Mycobacterium tuberculosis* WGS data. We then evaluate three approaches for reconstructing the underlying mixed constituent strain sequences. This allows these samples to be included in downstream analysis to gain insights into the epidemiology and transmission of mixed infections.

**Methods:** We employed a Gaussian mixture model to cluster allele frequencies at mixed sites (hSNPs) in each sample to identify signals of multiple strains. Building upon our previous tool, MixInfect, we increased the accuracy of classifying *in vitro* mixed samples through multiple improvements to the bioinformatic pipeline. Major and minor proportion constituent strains were reconstructed using three approaches and assessed by comparing the estimated sequence to the known constituent strain sequence. Lastly, mixed infections in a real-world *Mycobacterium tuberculosis* population from Moldova were detected with MixInfect2 and clusters of recent transmission that included major and minor constituent strains were built.

**Results:** All 36/36 *in vitro* mixed and 12/12 non-mixed samples were correctly classified with MixInfect2, and major strain proportions estimated with high accuracy, outperforming previous tools. Reconstructed major strain sequences closely matched the true constituent sequence by taking the allele at the highest frequency at hSNPs, while the best performing approach to reconstruct the minor proportion strain sequence was identifying the closest non-mixed isolate in the same population, though no approach was effective when the minor strain proportion was at 5%. Finally, fewer mixed infections were identified in Moldova than previous estimates (6.6% vs 17.4%) and we found multiple instances where the constituent strains of mixed samples were present in transmission clusters.

**Conclusions:** MixInfect2 accurately detects samples with evidence of mixed infection from WGS data and provides an excellent estimate of the mixture proportions. While there are limitations in reconstructing the constituent strain sequences of mixed samples, we present recommendations for the best approach to include these isolates in further analyses.

## Background

Mixed *Mycobacterium tuberculosis* (Mtb) infections occur when multiple, distinct strains of the pathogen are present simultaneously in a single host ^1^. These complex infections can be common and have been estimated to occur in upwards of 20% of patients with culture-positive tuberculosis (TB), particularly in high-burden settings with high-levels of exposure to infectious individuals ^2–7^. Characterizing the full pathogen strain diversity within individuals can have important implications for patient-level outcomes (e.g., hetero-resistance) and for accurate transmission inference ^1,8–10^.

Whole genome sequencing (WGS) data can be used to identify mixed *Mtb* infection by detecting genomic loci with evidence of more than one allele. A high proportion of these heterozygous single nucleotide polymorphisms (hSNPs) can be indicative of multi-strain infection ^6,11^, while some may represent within-host clonal heterogeneity or random sequencing errors. However, in most standard analysis pipelines, samples with a high number of hSNPs are often either removed from further analysis to account for potential mixed infections, or a single consensus sequence is produced for each isolate. In the latter case, the final sequence is composed of consensus SNPs (cSNPs) with only one nucleotide supported at a locus; when hSNPs occur, the nucleotide with the highest allele frequency above a given threshold is used ^12,13^. Consequently, the sequence may contain an excess of ambiguous base calls where the major allele frequency at hSNPs is below the chosen threshold to confidently assign a consensus nucleotide.

Failing to account for mixed infection can impact the accuracy of phylogenetic trees and subsequent phylodynamic analysis, as well as transmission reconstructions supported by genomic data. When a single consensus sequence is used for mixed samples, potential variation that would increase the genomic divergence between hosts may be ignored, incorrectly linking individuals in transmission chains ^8,11,14^. Furthermore, true transmission events may be rejected when the transmission of a minority strain from an infected individual occurs ^15^. When isolates with excess hSNPs are removed from the analysis entirely, they will necessarily be missing from all inferred transmission pairs, whose accuracy may therefore suffer. To accurately reconstruct transmission networks that include hosts with mixed infection, constituent strains must be delineated from the sequencing data and the distinct consensus sequences reconstructed and included in onward analysis.

Previous methods have been developed that detect mixed infections from microbial WGS data. These approaches work either by using probabilistic models to identify clusters in the sequencing reads or hSNP allele frequencies in assemblies ^6,9,16^, or by computing the likelihood of more than one sequence being present in the sequence data by matching to a reference database ^11,17^. Some of these methods also estimate the constituent sequences in mixes by grouping reads or alleles based on cluster designation ^16^ or by assigning the constituent strains as the closest matched strain in a database, which requires the constituent strain to be present in this database ^17^. A key limitation for determining the accuracy of these approaches has been the absence of real-world mixed samples to use as a gold standard for detecting mixtures and their proportions. Typically, these tools have been evaluated using synthetic sequences that may not reflect relevant properties of sequence data from mixed infections.

WGS data are available for 36 two-strain TB samples that were produced *in vitro* from *Mtb* isolates collected in Malawi ^18,19^ in different majority and minor strain proportions ^6^. These samples likely represent a closer approximation to true mixed TB infection and can be used to evaluate approaches for identifying multi-strain samples and reconstructing their constituent strains. Here, we present a new tool for identifying multi-strain Mtb samples from WGS data, **MixInfect2,** that builds upon a previously a published method ^6^ to improve the accuracy of mixed infection detection. We compare this new method to three previous tools for identifying mixed infection (MixInfect ^6^, SplitStrains ^16^, and QuantTB ^17^) and then assess different approaches for reconstructing the constituent strain sequences of the 36 *in vitro* mixed samples at varying major and minor strain proportions. Finally, we apply the optimal approach to a real-world dataset of over 2,000 *Mtb* isolates collected from the Republic of Moldova between 2018 - 2019 to predict the constituent strain sequences of mixed samples and place these strains into putative transmission clusters. This work allows us to gain insights into the epidemiology and transmission of mixed *Mtb* infection.

## Methods

### Sample data and genomic sequence analysis

WGS data was obtained for 36 *in vitro* mixed Mtb samples and their 12 non-mixed constituent strains (ENA project ID PRJEB2794). Mixed samples were produced using Mtb DNA obtained from 12 TB culture positive patients that were part of a larger cohort of 2,056 isolates from the Karonga district of Malawi between 1995 and 2014, for which WGS data is also available. To create the mixed samples, known concentrations of DNA from two isolates were combined at appropriate volumes *in vitro* to produce samples with major and minor strain proportions of 70/30, 90/10, and 95/5. The 12 constituent strains were also sequenced without mixing to represent ‘pure’ strains. These samples also represented both within and between major MTBC lineage mixes. Full details of the sample preparation and Illumina short-read sequencing of both the *in vitro* mixed samples and all Karonga strains are available at Sobkowiak *et. al*., 2018 ^6^.

We used an Illumina short-read WGS dataset of 2,220 Mtb isolates collected between 2018 and 2019 in the Republic of Moldova to investigate the prevalence and potential transmission of mixed infection in this population. These isolates represent around 80% of the 2,770 individuals notified with culture-positive TB in that period ^20^. Further details on the sample data and sequencing of Moldova isolates are available at Yang *et. al.,* 2022 ^21^.

For each isolate, WGS data were inspected using FASTQC software and aligned to the H37Rv reference strain (NC_000962.3) using the BWA-MEM algorithm ^22^. BAM files were created and sorted using SAMtools ^23^ and the final alignment files assessed for mapping quality using the SAMtools ‘flagstat’ command. Variant calling was carried out using the GATK ‘HaplotypeCaller’ and ‘GenotypeGVCFs’ commands ^24^, with joint calling for all isolates in each of the Karonga and Moldova datasets carried out separately. SNPs were removed if they failed quality control using the following filters with the ‘VariantFiltration’ option in GATK: QD < 2.0, QUAL < 30.0, SOR > 3.0, FS > 60.0, MQ < 40.0. Finally, SNPs identified in *pe/ppe* genes, known antimicrobial resistance genes, and repetitive regions were removed (**Supplementary file**).

### Detecting mixed infection and estimating proportions

Our new method to identify mixed infection from WGS data extends a previously developed tool, **MixInfect** ^6^, that uses a Gaussian mixture model (GMM) to cluster the allele frequencies at hSNPs and determine the number of groups in the data with the highest likelihood. Where a model with two or more groups is selected based on the Bayesian information criterion (BIC) value, a sample is declared to be a ‘mix’, otherwise it is inferred to be ‘pure’ and contain only one strain. In predicted mixed samples, the proportion of majority and minority strains is estimated by taking the mean allele frequencies of sites assigned to each group. We have made four key modifications, enumerated below, that improve the accuracy of detecting mixed samples by reducing the number of false positive SNP calls. We call our new tool **MixInfect2**.

1. The variant call file (VCF), built from either a single sample or multiple samples, is used to determine cSNPs and hSNPs rather than the per-sample alignment (BAM) file. This allows for fewer false positive SNP calls as the filtration steps in variant calling are applied.
2. hSNPs found within a 100bp window are combined and the median major and minor allele frequencies taken as a single data point. This will minimize the impact of regions of potential alignment or sequencing error, which will reduce the number of false positive hSNPs.
3. hSNPs are classified directly from allele frequencies per site given a minimum read depth of the minority variant of 10 reads rather than relying on imposing a diploid genome option to determine mixed sites in variant calling.
4. For population-level analysis rather than single samples, hSNPs found at a higher frequency than a given threshold across all samples, including ‘pure’ strains, can be masked. This option removes hSNPs that are present at high frequencies in the dataset due to systematic sequencing or alignment issues. It also removes hSNPs that may be present due to higher levels of within-host variation, such as from loci under selection, rather than through the presence of multiple, distinct strains from mixed infection.

### Reconstructing constituent strains of mixed infection

Different approaches have been used previously to reconstruct full constituent sequences or assign specific variants to different strains in mixed TB infections. These fall into two broad categories; either constituent sequences are inferred by assigning sequencing reads or nucleotides at mixed sites to major and minor strain sequences using allele frequencies ^9,16^ or by considering constituent strain sequences as the closest matching strain from a reference database ^17,25^. Here, we tested the following three approaches for predicting the major and minor constituent sequences for each *in vitro* mixed sample and calculated the SNP distance between the final inferred sequence and the true constituent strain sequence.

1. ‘**Consensus allele frequency’** – The highest frequency allele (> 50%) at hSNPs was assigned to the major strain and lowest frequency allele (< 50%) assigned to the minor strain. Where allele frequencies were equal, an ambiguous call ‘N’ was assigned. All invariant sites were called as the reference nucleotide and all cSNPs called as the respective alternative nucleotide in each sequence per sample.
2. ‘**Closest strain’** – The constituent strains of mixed samples were predicted to be the sequence of the closest ‘pure’ (non-mixed) strain (by SNP distance) found in the reference database, which comprised all non-mixed samples in the population. The pairwise SNP distance was calculated between each pure strain sequence and the mixed sample at both cSNPs and hSNPs. In hSNPs, if any allele supported by 10 reads at that site in the mixed sample was present in the pure strain, the distance at that position was 0. The major constituent strain sequence was estimated to be the sequence of the ‘pure’ strain in the database in which the majority of alleles at hSNPs of the mixed sample were the highest allele frequency (> 50%), then which had the minimum SNP distance to the mixed sample, and then had the lowest proportion of ambiguous base calls. The minor constituent strain sequence was predicted in the same way but chosen from the ‘pure’ strains in the database in which the majority of alleles at hSNPs of the mixed sample were found at the lowest allele frequency (< 50%). If more than one ‘pure’ strain met these criteria as either the major or minor constituent strain, any allele differences between the chosen ‘pure’ strains were called either as the cSNP allele from the mixed sample variant call or given an ambiguous base call, “N”, at hSNPs in the final reconstructed sequence. If no ‘pure’ strains were present in the database that had the majority of alleles at the hSNPs of the mixed sample at either the high or low allele frequency, a constituent strain sequence was not predicted.
3. ‘**Closest strain + SNPs’** – The closest major and minor strain in the reference database was found using the ‘closest strain’ method described above. The allele at hSNPs in major and minor constituent sequence was called as the nucleotide at this position in the closest non-mixed strain. For the sequence at other sites, invariant sites were called as the reference nucleotide and all cSNPs in the mixed sample called as the alternative nucleotide.

All scripts to run **MixInfect2,** including identifying mixed infections and reconstructing constituent strains, can be found at github.com/bensobkowiak/MixInfect2.

### Transmission analysis of Moldovan M. tuberculosis isolates

Mixed infections were detected in the 2,220 Mtb isolates collected in Moldova and the constituent strains predicted using the best performing approach identified from the analysis of *in vitro* mixed samples. Lineage calling and antimicrobial resistance profiling of the constituent sequences was carried out by detecting the presence of lineage-specific SNPs and rifampin and isoniazid resistance-conferring mutations in a database constructed from SNP sets contained in TB-profiler ^26^. Putative clusters of recent transmission were produced by linking sequences in the same group with a pairwise distance of ≤ 5 SNPs, which has been used previously to identify recent transmission ^27^. All statistical analyses were performed in R.

## Results

### Detecting mixed infection using in vitro mixed samples

We compared the accuracy of our new tool, **MixInfect2,** against previous methods (**MixInfect** ^6^, **SplitStrains** ^16^ and **QuantTB** ^17^) for detecting mixed infections and to estimate the major strain proportion from the dataset of 36 *in vitro* mixed samples and 12 non-mixed (‘pure’) strains. The average coverage in all these samples was relatively high, ranging from 356- to 482-fold. We found that **MixInfect2** accurately classified 36/36 mixed samples as combinations of two strains and all pure samples as single strains in the dataset (**Figure 1**). In comparison, **QuantTB** identified four mixed samples as non-mixed (two at 90/10 and two at 95/5 mixing proportions) and two pure samples as mixes, as well as incorrectly predicting that six mixed samples were comprised of three strains. **SplitStrains** software correctly classified all mixed samples, but all pure samples were also predicted to be mixed strains and **MixInfect** incorrectly predicted one 90/10 and eight 95/5 mixed samples as pure strains.

**Figure 1.**
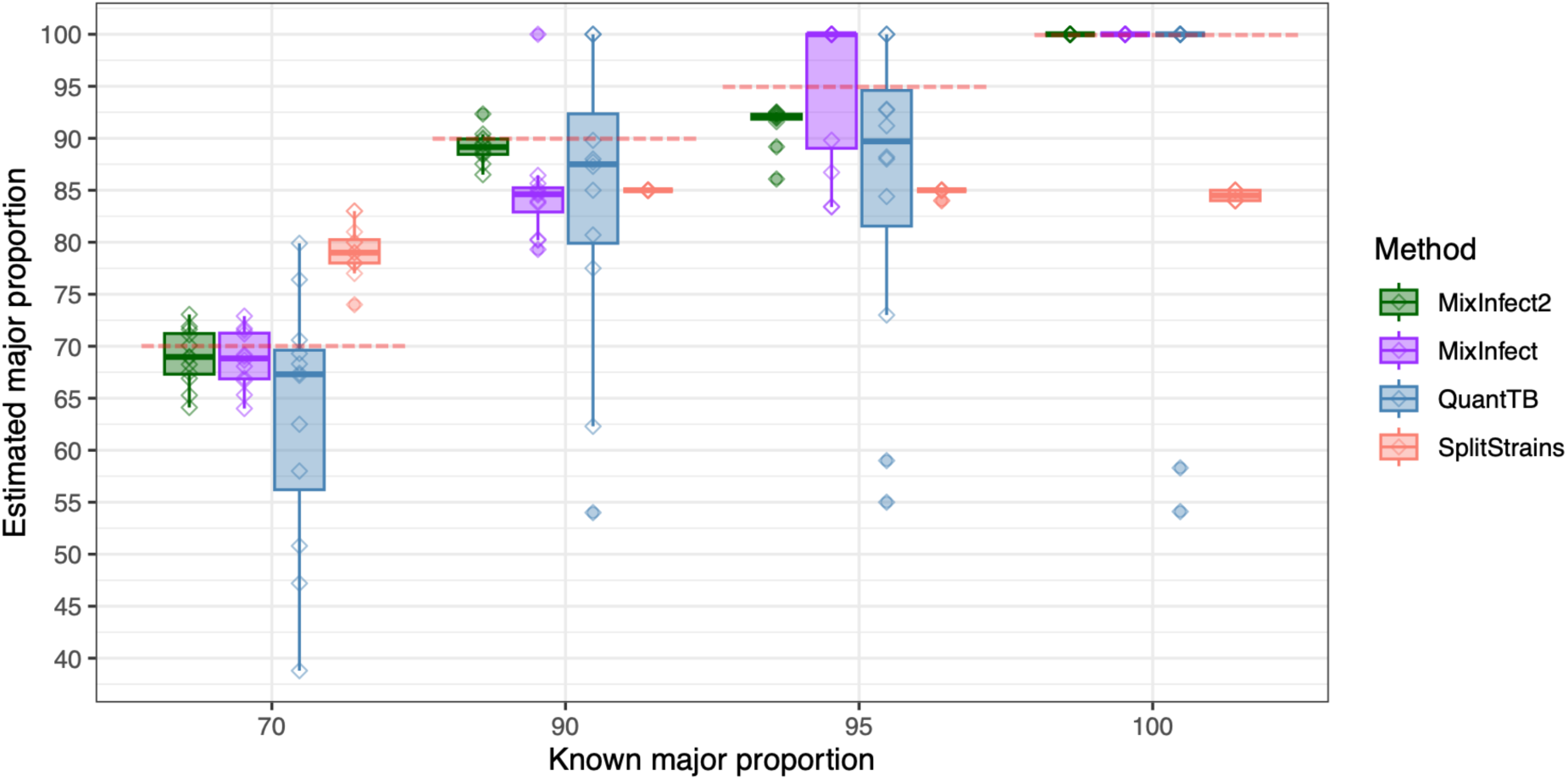
The estimated major strain proportion against the known major strain proportion of the 36 *in vitro* mixed samples and 12 single strain samples estimated using MixInfect2, MixInfect, SplitStrains, and QuantTB. Dashed red lines represent the true major strain proportion.

Furthermore, the estimated major strain proportion of all *in vitro* mixed samples was close to the known major strain proportion using **MixInfect2** (**Figure 1**), and our new approach significantly outperformed the other methods overall. In 70/30 major/minor strain proportion mixes, the median major strain proportion predicted by **MixInfect2** was 69.0% (IQR 67.3 – 71.2), with the absolute difference between the predicted and known proportion significantly different to estimates from both **QuantTB** and **SplitStrains** (t-test P < 0.05). In 90/10 mixes, the median major strain proportion predicted by **MixInfect2** was 89.2% (IQR 88.5 – 89.9), with the predictions significantly different to **QuantTB** and **MixInfect** (t-test P < 0.05). Finally, in 95/05 mixes the median major strain proportion was estimated as 92.1% (IQR 91.8 – 92.3) and this was significantly different to **QuantTB** and **SplitStrains** (t-test P < 0.05).

### *Reconstructing constituent strain sequences of* in vitro *mixed samples*

We next compared three approaches for reconstructing both the major and minor constituent strain sequences of *in vitro* mixed samples as detailed in the methods section: 1) consensus allele frequency, 2) closest strain, and 3) closest strain + SNPs. For approaches that used a reference dataset to find the closest strain in non-mixed samples from the population, we included the sequences from a larger cohort of 2,056 TB culture-positive individuals in the Karonga District of Malawi from which the constituent strains of the *in vitro* mixed samples were obtained. Of these isolates, 80 assemblies failed quality control and 189 samples were identified as mixed infection using **MixInfect2** and were removed, along with the 12 strains that matched the pure strains in the *in vitro* mixed dataset to avoid replication of these strains in the database. This resulted in a final reference database of 1775 non-mixed clinical strains and the 12 pure strains from the *in vitro* dataset.

### Reconstructing major strain sequences

Figure 2 shows the median SNP distance between the inferred major strain sequence and known constituent sequence for the 36 *in vitro* mixed samples predicted using the three approaches for reconstructing mixed sequences. We first tested the three approaches when the constituent strains of mixed samples (the pure strains from the *in vitro* dataset) were included in reference dataset (Figure 2A). The inferred sequence of the major strains was very close to the true sequence when estimated by all approaches (Figure 2A), with all sequences predicted to within 5 SNPs of the known sequence. The median SNP distance between the predicted sequence and true sequence was 0 SNPs for all methods, apart from the ‘closest strain + SNPs’ method at 95% major strain proportion, which was 1 SNP, and there was no significant difference between the tested approaches (Kruskal-Wallis p > 0.05).

**Figure 2.**
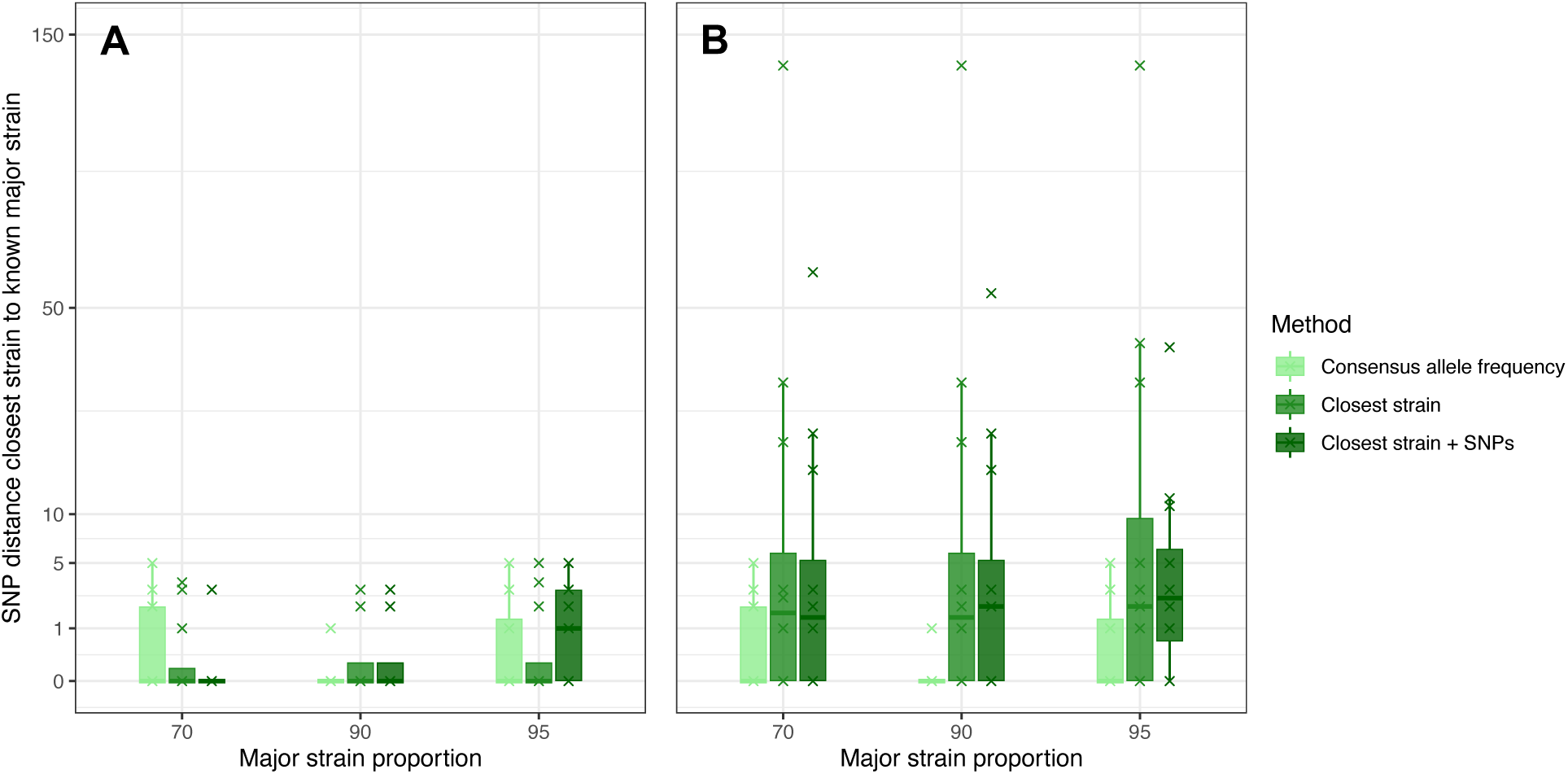
Boxplots showing the SNP distance between the predicted and known major constituent strains of *in vitro* mixed samples using the three tested approaches. Boxes are colored by the different approach used to predict the constituent strain sequence. Plot A shows the results when the known constituent sequences are included in the reference database, and plot B shows the results when these sequences are removed. Note that the vertical axis has been transformed by the square root for visualization.

When the constituent strains of the mixed samples were not included in the reference dataset, the ‘closest strain’ and ‘closest strain + SNPs’ methods performed slightly worse than when consensus strains were included (Figure 2B). Although there was still no significant difference among the tested methods when comparing the median SNP distance between the inferred and known sequences (Kruskal-Wallis p > 0.05), the average SNP distance was larger using these two methods when constituent sequences were removed from the database. Compared to the results when including the known constituent ‘pure’ strain in the database, the median SNP distance between the inferred sequence and true constituent strain sequence increased to 1.5 – 2.5 SNPs, with a maximum of 136 SNPs difference. As the ‘consensus allele frequency’ approach does not use a reference database, predicted sequences did not change with this approach when removing the constituent strains, resulting in the same median of 0 SNP distance and maximum of 5 SNP distance between predicted and known sequences. As such, the ‘consensus allele frequency’ appears to offer the best approach to reconstruct the majority strain sequence of mixed infection.

### Reconstructing minor strain sequences

The performance of the tested approaches for reconstructing the minor strain sequences of the *in vitro* mixed samples was impacted more by the proportion of the minor strain in the mix than for the major constituent strain. At the 30% minor strain proportion, the ‘closest strain’ method inferred minor strain sequences that most closely matched known constituent sequences, with a median of 0 SNPs and a maximum of 3 SNPs difference (Figure 3A). The ‘closest strain + SNPs’ method also found closely linked sequences with a median 0 SNP distance between predicted and known sequences but with a higher maximum distance of 7 SNPs. The ‘consensus allele frequency’ method performed significantly worse (Kruskal-Wallis p < 0.05), with a median of 5.5 SNPs between the predicted and known constituent sequence. Removing the true constituent sequences from the dataset when searching for the closest strains increased the median SNP distance of the ‘closest strain’ and ‘closest strain + SNPs’ methods to 2 SNPs (Figure 3B). This was still lower than the ‘consensus allele frequency’ approach but difference between the tested methods was now not significant (Kruskal-Wallis p > 0.05).

**Figure 3.**
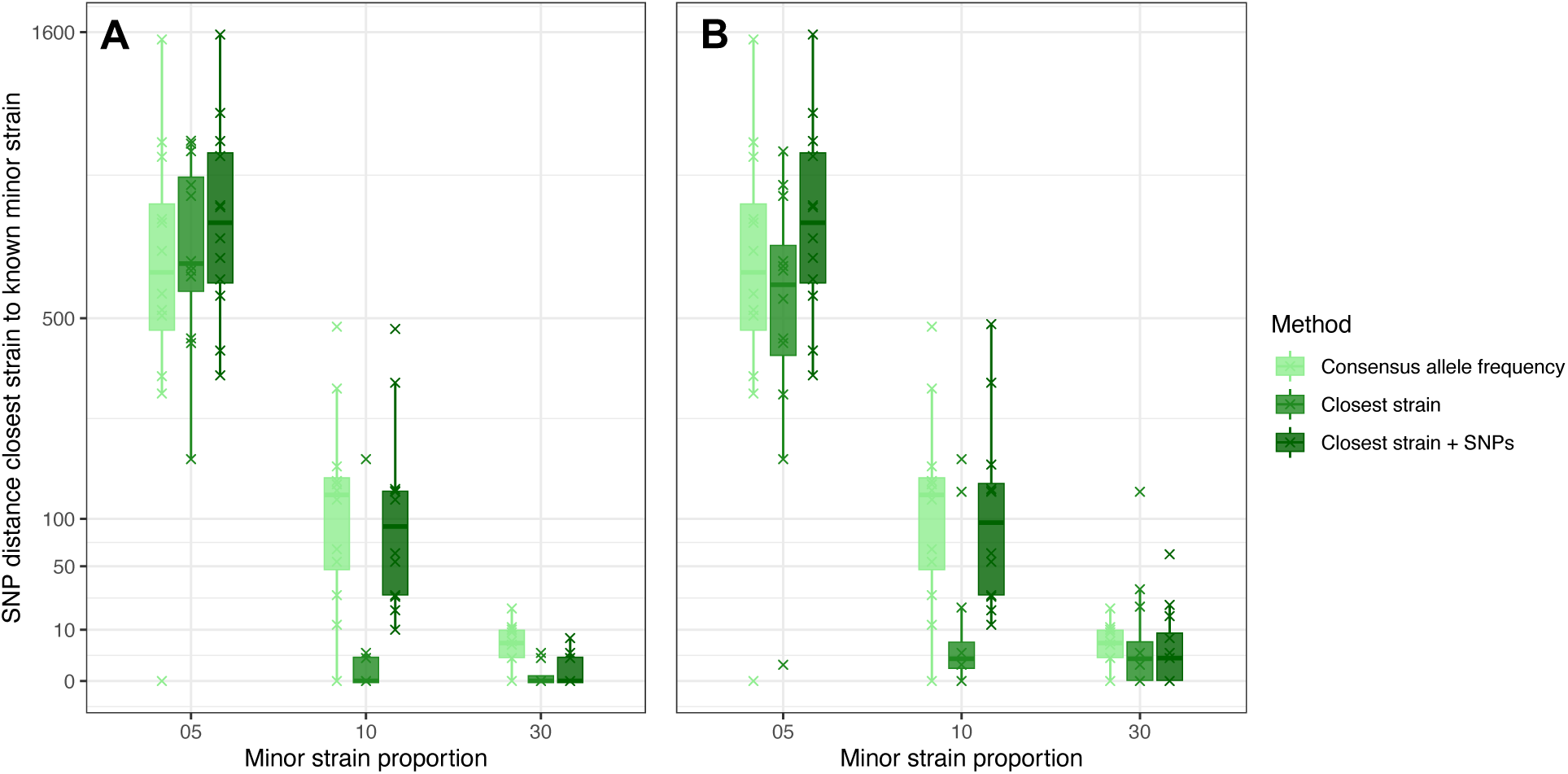
Boxplots showing the SNP distance between the predicted and known minor constituent strains of *in vitro* mixed samples using the three tested approaches. Boxes are colored by the different approaches used to predict the constituent strains. Plot A shows the results when the known constituent strains are included in the list of ‘pure’ strains, and plot B shows the results when these strains are removed. Note that the Y axis has been transformed by the square root for visualization.

When the minor strain proportion was at 10% in the *in vitro* mixed samples, the ‘closest strain’ approach performed significantly better than other methods (Kruskal-Wallis p < 0.05), with a median SNP distance between the predicted and known constituent sequence of 0 SNPs. This compared to 131.5 SNPs using the ‘consensus allele frequency’ method and 93.5 SNPs using the ‘closest strain + SNPs’ method (Figure 3A). While there was one outlier sequence with a large SNP distance between predicted and true sequence using the ‘closest strain’ approach (maximum 187 SNPs), all other sequences were predicted within 3 SNPs of the true sequence. In addition, when the known constituent sequence was removed from the dataset, this method still outperformed the other approaches significantly and the median SNP distance between predicted and true sequences was 2 SNPs (Figure 3B).

At a 5% minor strain proportion in mixed samples, the median SNP distance between the predicted sequence and known constituent strain sequence was high using all tested approaches (Figure 3). Inspection of VCF file showed that many sites that differed between the minor and major constituent strains in the 95/5 samples were called as a cSNP matching only the allele of the major strain sequence instead of hSNPs. This was due to the very low number or absence of reads carrying the allele from minor constituent strain. This was also evidenced by the large difference in the SNP distance between the closest strain identified in the reference database and the known constituent strain in most of the *in vitro* minor strains at a 5% mixing proportion (**Supplementary figure S1**). In these samples, the closest strain identified was often very divergent from the known constituent strain and in some instances was closer to the major constituent strain, which was a different MTBC major lineage in some of the mixed samples. Thus, it appears that a minor strain proportion of 5% is too low to accurately infer the minor strain sequence using the WGS data and approaches considered here.

### Sensitivity analysis

To assess how the size and composition of the dataset affected the performance ‘closest strain’ and ‘closest strain + SNPs’ methods, we downsampled the dataset of ‘pure’ isolates in the Karonga dataset to 50% and 75% of the original size and re- calculated the SNP distance between the predicted and known major and minor sequences for all *in vitro* mixed samples. This process was repeated 100 times at 50% and 75%, randomly selecting ‘pure’ strain sequences to include in the new dataset. We found that the median SNP distance of the ‘closest strain’ method increased from 0 SNPs at all major strain proportions to 3 SNPs with 50% downsampled dataset (**Supplementary figure S2A**) and 4 SNPs with the 75% downsampled dataset (**Supplementary figure S2C**). The median SNP distance between predicted and known major strain sequences increased from 0 SNPs using the ‘closest strain + SNP’ approach to 2 – 5 SNPs in the 50% and 4 – 5 SNPs in the 75% downsampled dataset. Furthermore, the maximum distance to the true sequences when using these approaches increased dramatically in the downsampled datasets where sequences that were closely related to the major strain in the mixed sample may not have been included. As such, using the ‘consensus allele frequency’ approach appears to best the option to predict major constituent strain sequences to mitigate the possibility that close ‘pure’ strains are not included in the dataset.

For the minor strain sequence prediction, the optimal method to reconstruct the sequence appeared to depend more on the completeness of sampling and sequencing of the infected population. While the ‘closest strain’ approach performed the best in the 30% minor strain proportion samples with the full dataset, when the dataset is downsampled by 50% and 75% ‘consensus allele frequency’ achieves only a slightly higher median SNP distance between predicted and known sequences than the other approaches (5.5 SNPs compared to 2 – 4 SNPs) but with far fewer samples with a predicted sequence a large distance from the true constituent sequence (**Supplementary figures S2B** and **S2D**). At 10% minor strain proportion, the ‘closest strain’ approach still performs best for estimating the minor strain in both the 50% and 75% downsampled datasets but again the number of predicted sequences that were a large SNP distance to the known sequence was high. Therefore, in populations where sampling or sequencing is sparse and the likelihood of including the constituent strain or a closely related sequence is low it may be problematic to predict minor strain sequences, particularly for at low minor strain proportions. In this instance, it may be optimal to use the ‘consensus allele frequency’ method to predict both major and minor constituent sequences but not attempt to reconstruct minor strains at low mixing proportions to reduce the chances of poorly predicting the minor strain sequence.

In our bioinformatic pipeline, hSNPs were defined as sites with more than one allele supported in aligned reads and a minimum minor allele depth of 10 reads. We assessed the impact characterizing hSNPs in this way by comparing the SNP distance between the predicted sequence and the true constituent strain sequence using the ‘closest strain’ approach when changing the metrics used to call hSNPs. These included lowering the minimum minor allele read depth to 5 reads, using a minimum allele read proportion (rather than raw depth) of 0.01, 0.02, 0.05 and 0.1, and using the heterozygous base call (e.g., “0/1”) by setting the ploidy option the variant-calling software to diploid. We found no significant change in the results of reconstructing the major and minor constituent strains of the *in vitro* mixes using the ‘closest strain’ approach using the different hSNPs calling methods, apart from an increase in the distance between the predicted and known sequence at the 10% minor strain proportion when using a minimum minor allele proportion of 0.1 (**Supplementary figure S3**).

### Mixed TB infection and transmission in Moldova

We used a real-world dataset of 2220 *Mtb* isolates from the Republic of Moldova ^21^ to identify the proportion of mixed infection in this population and reconstruct the constituent strain sequences. A total of 146 of 2220 (6.6%) isolates were identified as mixed infection using **MixInfect2** (**Supplementary file**), substantially fewer than previously predicted in this dataset using the earlier **MixInfect** approach (386/2220; 17.4%) ^21^. All major constituent strain sequences were predicted using the ‘consensus allele frequency’ approach as this achieved the best results in the *in vitro* mixed samples. The ‘closest strain’ approach was used to predict minor strain sequences with an estimated minor strain proportion of ≥ 10%. With this approach, we also set a maximum distance threshold of 1000 SNPs to the closest strain in the reference database to reduce the chance of matching a strain that is very divergent to the true constituent strain; if the closest strain was further then no minor strain sequence was predicted. We did not reconstruct the major or minor strain in mixed samples with a major strain proportion estimate of ≥ 50% and < 60% (N = 15) as we have not reliably tested these methods when constituent strain proportions were close to parity. Eleven isolates with a high proportion of hSNPs (≥ 10% of total SNPs) that were not classified as mixed infection were flagged and removed from further analysis. This removal resulted in a final dataset of 2291 isolates: 2063 ‘pure’ strain, 129 major strain, and 99 minor strain sequences.

In the 99 mixed samples for which both the major and minor strain sequence was predicted, we found that 45 (45.4%) contained a mix between different major lineages and a further 23 (23.2%) were mixes of lineage 4 sub-lineages (all lineage 2 strains were of the Beijing lineage 2.2). We also found evidence of hetero-resistance to isoniazid and/or rifampin in 27/99 mixed infections (27.3%), and while this can occur in single strain infections when these SNPs can be under selection and not reached fixation, this proportion was higher than hetero-resistance found in 20/2063 non-mixed strains (Chi-square test 295.08, P < 0.05). A maximum likelihood phylogeny that included both non-mixed and the predicted constituent mixed strain sequences showed that most mixed constituent strains were closely related to other sequences in the dataset, although there were a small number of major strains that appeared relatively genetically distant to any other strain in the dataset (**Supplementary figure S4**).

Finally, transmission clusters were constructed by linking all sequences that were separated by a pairwise SNP distance of ≤ 5 SNPs, including the predicted major and minor mixed constituent strains. We identified 90 clusters that contained at least one constituent strain of the mixed infections, including a large transmission cluster containing 130 sequences with six major constituent and nine minor constituent strains of mixed infections, and a large cluster of 66 sequences containing two major constituent and one minor constituent (Figure 4). A total of 45 of 129 (34.9%) major constituent strains were predicted to be part of a transmission cluster compared to 951/2063 (46.1%) of pure strains (Chi-square test 5.71, P = 0.02). A total of 96 of 99 minor constituent strains were found in transmission clusters, although this high number of minor strains included in clusters was due to the ‘closest strain’ method predicting the minor constituent sequence to be the near identical to the sequence of the closest ‘pure’ strain in most instances. There also appeared to be one cluster comprising only one non-mixed strain that was the closest strain sequence to 14 minor strains. While it could be possible that this strain is closely related to the minor strain in these mixed samples, it may be that this is instead evidence of minor contamination in these samples rather than clinical mixed infection. The full list of sequences with cluster designation and lineage can be found in **Supplementary file**.

**Figure 4.**
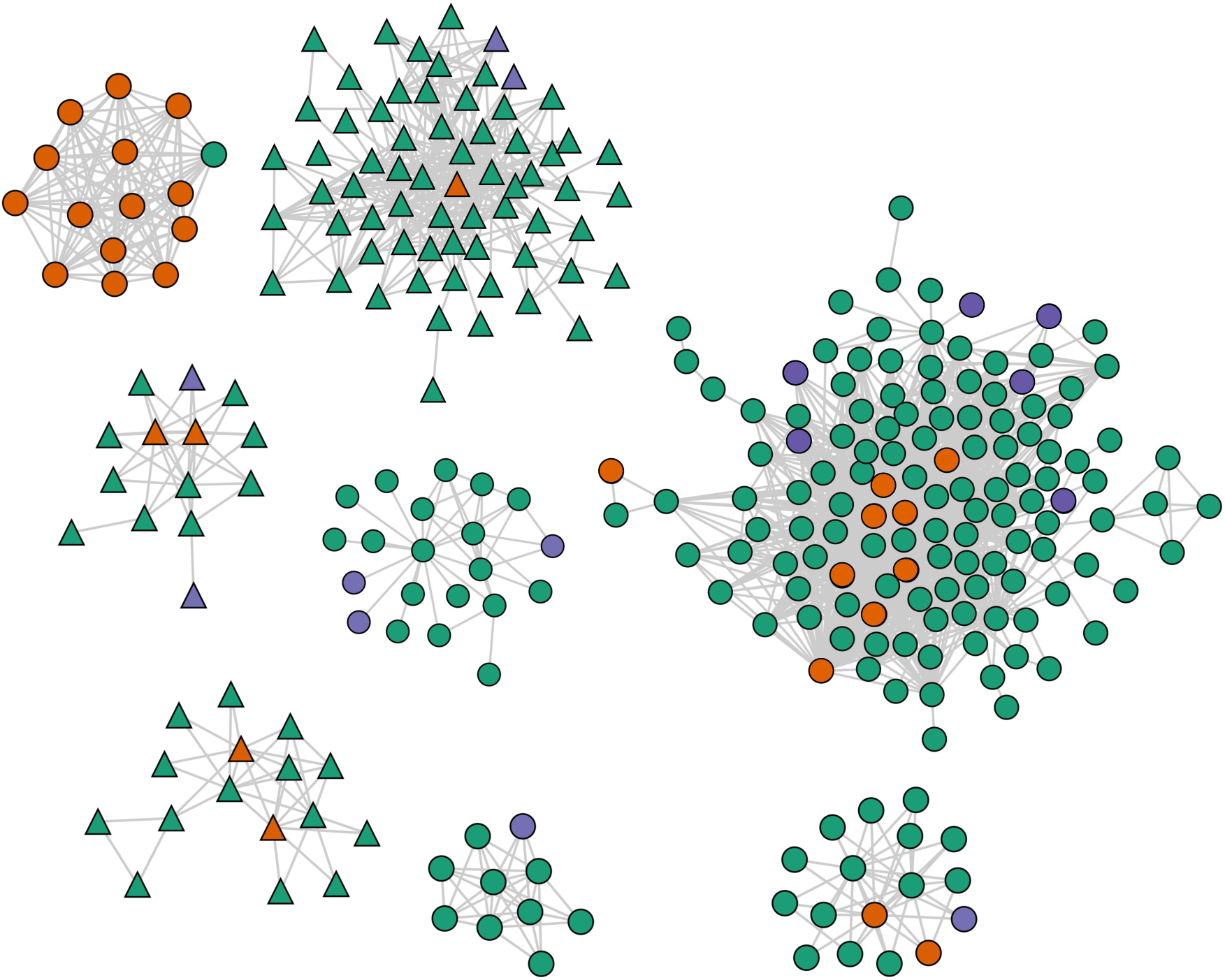
Visualization of the eight largest transmission clusters (N > 10) in the Moldova *Mtb* dataset that contained at least one mixed constituent strain produced using TGV (https://jodyphelan.github.io/tgv/). The color of the node represents whether the sample is non-mixed (green), the major constituent strain of mixed infection (blue), or minor constituent strain of mixed infection (red). Edges represent any pairwise distance between sequences of ≤ 5 SNPs and the node shape denotes the MTBC major lineage, where circle nodes are lineage 4 strains and triangle nodes lineage 2 strains.

## Discussion

Mixed microbial infections with two distinct strains present in a single host are not rare and can complicate population genomic analysis. Here, we have improved on the performance of previously developed tools to detect mixed TB infection from WGS data and evaluated different approaches to reconstruct the constituent strain sequences in mixed samples. Their application allows us to identify mixed infection more accurately in Mtb populations directly from WGS data and include two sequences representing major and minor strain sequences from these individuals in further analysis rather than either erroneously inferring a single sequence or removing these samples entirely. This work can also be applied to detecting mixes with other pathogens that can cause polyclonal infection, such as *Staphylococcus aureus* ^28^ and *Helicobacter pylori* ^29^.

Our new tool, MixInfect2, identified all mixed and non-mixed strains in a dataset of *in vitro* mixed samples and performed better at detecting these mixes from sequence data than three other methods (MixInfect, SplitStrains, and QuantTB). These *in vitro* samples were previously analyzed using SplitStrains ^16^ and MixInfect ^6^, though the results for the 12 non-mixed strains in the dataset were not reported in the SplitStrains study. In our analysis, SplitStrains predicted that all non-mixed samples were mixed infections, and MixInfect incorrectly classified nine mixed samples as pure. These tools also used allele frequencies at mixed loci in sequence alignments to identify mixes, but the additional steps implemented in MixInfect2 to filter poor quality hSNPs and reduce false positive calls from the variant call file improved the accuracy in classifying mixed samples. The *in vitro* samples were not previously analyzed with QuantTB, but we found that using a reference database to quantify the number of strains in each sample was not as accurate as MixInfect2, with some samples only matching one strain in the database and a number of samples predicted to have more than two strains.

With our improved method, we detected fewer mixed infections than previously estimated using MixInfect in the dataset of Mtb isolates collected in the Republic of Moldova ^21^. This difference was due to the removal of potentially false positive hSNPs by the additional filtering steps now applied in MixInfect2, notably combining hSNPs within a 100bp window to prevent multiple variants found in close proximity in the genome to be treated as independent observations. While there may be instances where hSNPs are close on the genome, this will be more common in regions with systematic mapping or assembly issues caused by sequence duplication or non-specific mapping of the reads. Indeed, some samples previously detected as mixed infection in these data that were now classified as non-mixed still retained evidence of a high number of hSNPs, but inspection of the raw alignment (BAM) files found that most of these hSNPs were found in genomic regions with higher-than-average coverage, likely due to mis-mapping (**Figure S5**).

We found that majority constituent strain sequences of the *in vitro* mixed samples could be predicted by combining nucleotides from consensus calls (cSNPs and reference calls) and the highest frequency alleles at mixed sites (hSNPs) detected in VCF files. These predicted sequences were found to be very close (within 5 SNPs) to the known majority constituent strain sequence. However, inferring the minor strain sequence of the *in vitro* mixed samples was more difficult and the accuracy of the predicted sequence depended on the minority strain mixing proportion. When the minor strain proportion was at 5%, many alleles found in the minor constituent strain were instead called only as the allele of the major strain and any SNP differences between the major and minor strains were not detected. When the constituent strains of mixes are closely related there will be a low number of sites that differ and thus a closer minor strain may be found even when there is a low minor strain proportion as fewer will be wrongly called, although actually detecting mixed infections in samples comprising very similar underlying strains will be more difficult. Given these issues, along with a potential loss of signal from low frequency variants in the sampling, sequencing or bioinformatic stages of analysis, it is likely that some mixed infections could be missed if strains are present in very low frequencies in the host or where constituent strains are very closely related.

While the ‘closest strain’ approach was found to estimate the minor constituent strain best for most mixed samples with a higher minor strain proportion (≥ 10%), we also showed that reducing the size of the reference database of ‘pure’ strains increased the distance between the inferred sequence and known constituent strain. This was due to the reduced likelihood of including a closely related strain to the minor strain in the smaller reference database. Therefore, it is important to ensure that the sampling density in a population is high in such analysis to increase the chance of capturing ‘pure’ strains that are close to the constituent strain.

Using the optimal methods identified for estimating the constituent strain sequences in mixed samples, we found evidence of recent transmission including hosts with mixed infection in Mtb isolates collected in the Republic of Moldova. The Moldova data includes sequences from a high proportion of the individuals diagnosed with TB in the country during the study period ^21^ and thus it is likely that closely related strains to the minor strain in mixed infections were present in the reference database. We note that more variation may actually be present in several of the minor strain sequences predicted here that would distance these sequences further from the closest strain; this could cause transmission inference based on the closest strain sequence to overestimate the proportion of minor strains that cluster. It could also be problematic to include these sequences in a full person-to-person transmission network reconstruction as there would be less variation between the inferred constituent sequence and the closest ‘pure’ strain sequence, potentially linking hosts incorrectly in direct transmission events when in reality they are further away in the transmission chain. That said, given our results from the Karonga *in vitro* mixed dataset which show that the ‘closest strain’ method predicts the true minor strain sequence closely in most instances, it is likely that a high number of the predicted minor strain sequences would still be found in transmission clusters.

In this study, we only tested short-read sequence data of between 100 – 150 base pairs as these data available for the *in vitro* mixed samples. Long-read sequencing may improve approaches to reconstruct constituent strains where a large number of variants from each strain in the mixed sample would be contained on the same read to reduce the uncertainty when reconstructing full genomes. Also, we did not explicitly test how variation in the reference sequence used or per-sample read depth impacted the ability to call mixed infection, although the *in vitro* mixed samples all had relatively high genomic coverage (median 462-fold, range 386- to 584-fold). It is likely that deeper sequencing allows us to detect mixed infections with smaller minor strain proportions as more variation from these strains will be detected in the sequence data. Similarly, sequencing directly from sputum may allow for more within-host variation to be captured in the sequence data in the absence of potential bottlenecks introduced from culturing ^13^, although recent research has disputed this assertion ^30^. Furthermore, evidence of more than one strain in a sample does not unequivocally show evidence of mixed infection within a host. Finding a high proportion of mixed samples with the same or very similar constituent strain sequences may be the result of lab contamination, particularly if the hosts carrying the strains are unlikely to have been in contact based on their epidemiology.

In future work, validating the results presented here using *in vitro* mixed samples that are composed of more than two strains and with constituent strains at a 50/50 mixing proportion would improve methods to detect mixed infection and to reconstruct the constituent strains in real populations. Additionally, we could explore the signal from other genomic variants in mixed samples in addition to SNPs, such as small insertions and deletions or structural variants. Nonetheless, the results presented here provide a method for identifying Mtb mixed samples from short-read WGS data that enables these complex infections to be more accurately detected than previously possible. We have also evaluated the methods to reconstruct the constituent strains of mixed infections, with recommendations for the approach to use that best predicts the true genomic sequence. Carrying out this analysis in a real TB population from Moldova showed that recent transmission can involve both the major and minor constituent strains, suggesting that individuals infected with multiple strains can transmit to others and should be included in transmission investigations.

## Supplementary figures

**Supplementary figure S1.**
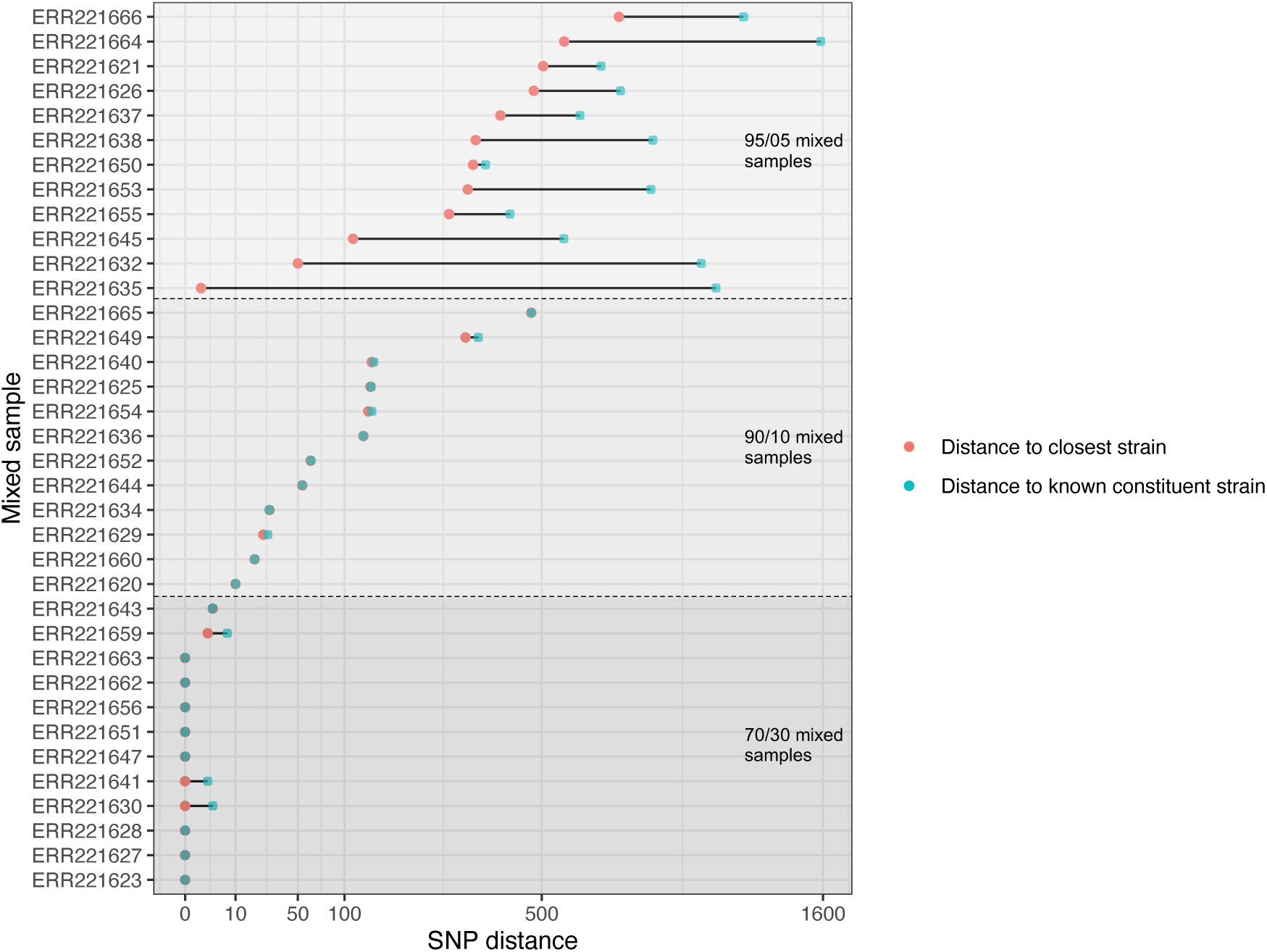
The SNP distance between the closest identified strain sequence in the reference database of ‘pure’ isolates in the Karonga dataset and the known constituent minor strain sequence in each *in vitro* mixed sample.

**Supplementary figure S2.**
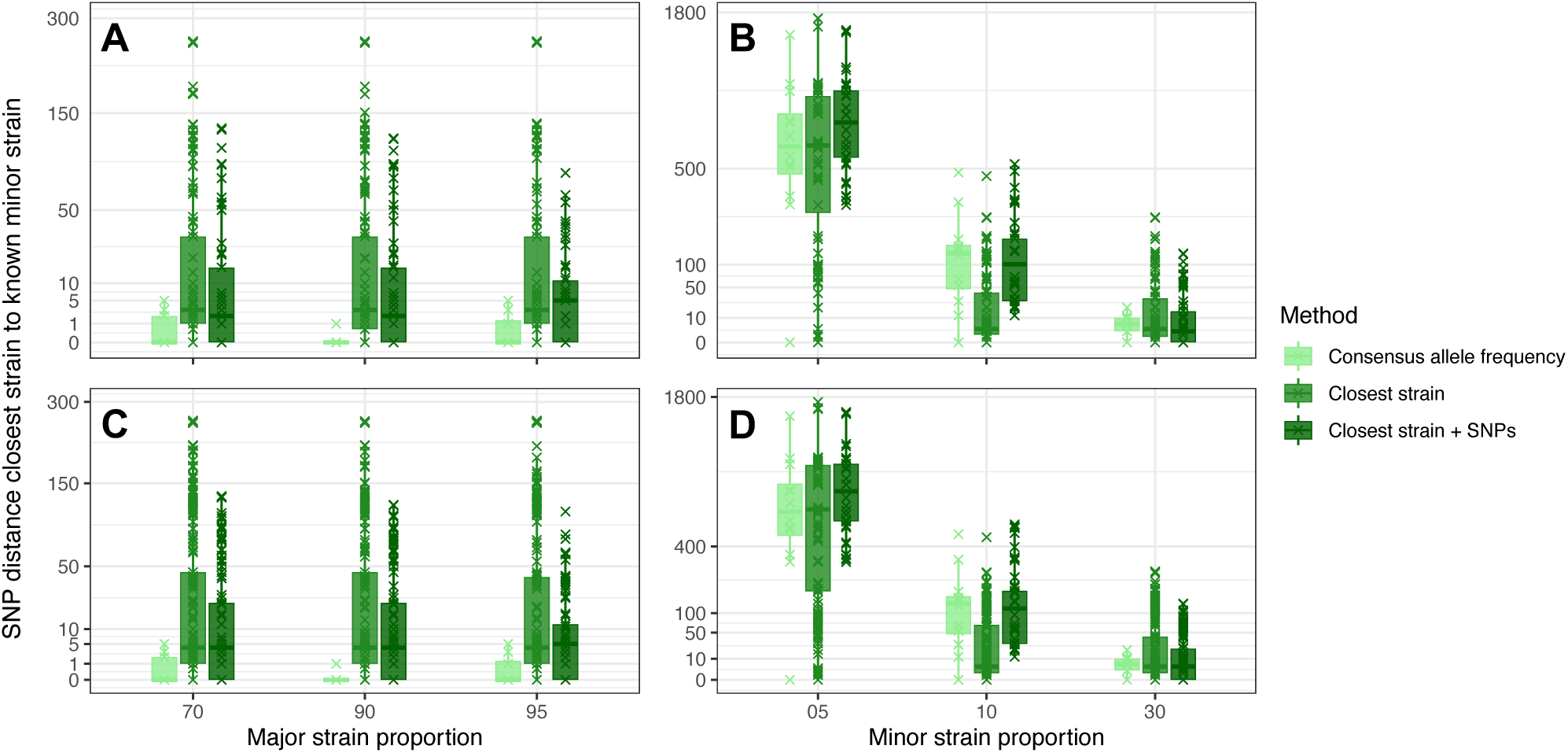
Boxplots showing the SNP distance between the predicted and known constituent strains of *in vitro* mixed samples using the three tested approaches when downsampling the reference dataset of ‘pure’ strains by 50% and 75%, with 100 replicates per downsample strategy. Plots A and B show the major and minor strain estimates with the 50% downsampled ‘pure’ dataset and plots C and D show the major and minor strain estimates with the 75% downsampled ‘pure’ dataset. Known constituent strains were not included in the ‘pure’ strain datasets. Boxes are colored by the different methods used to predict the constituent strains. Note that the Y axes have been transformed by the square root for visualization.

**Supplementary figure S3.**
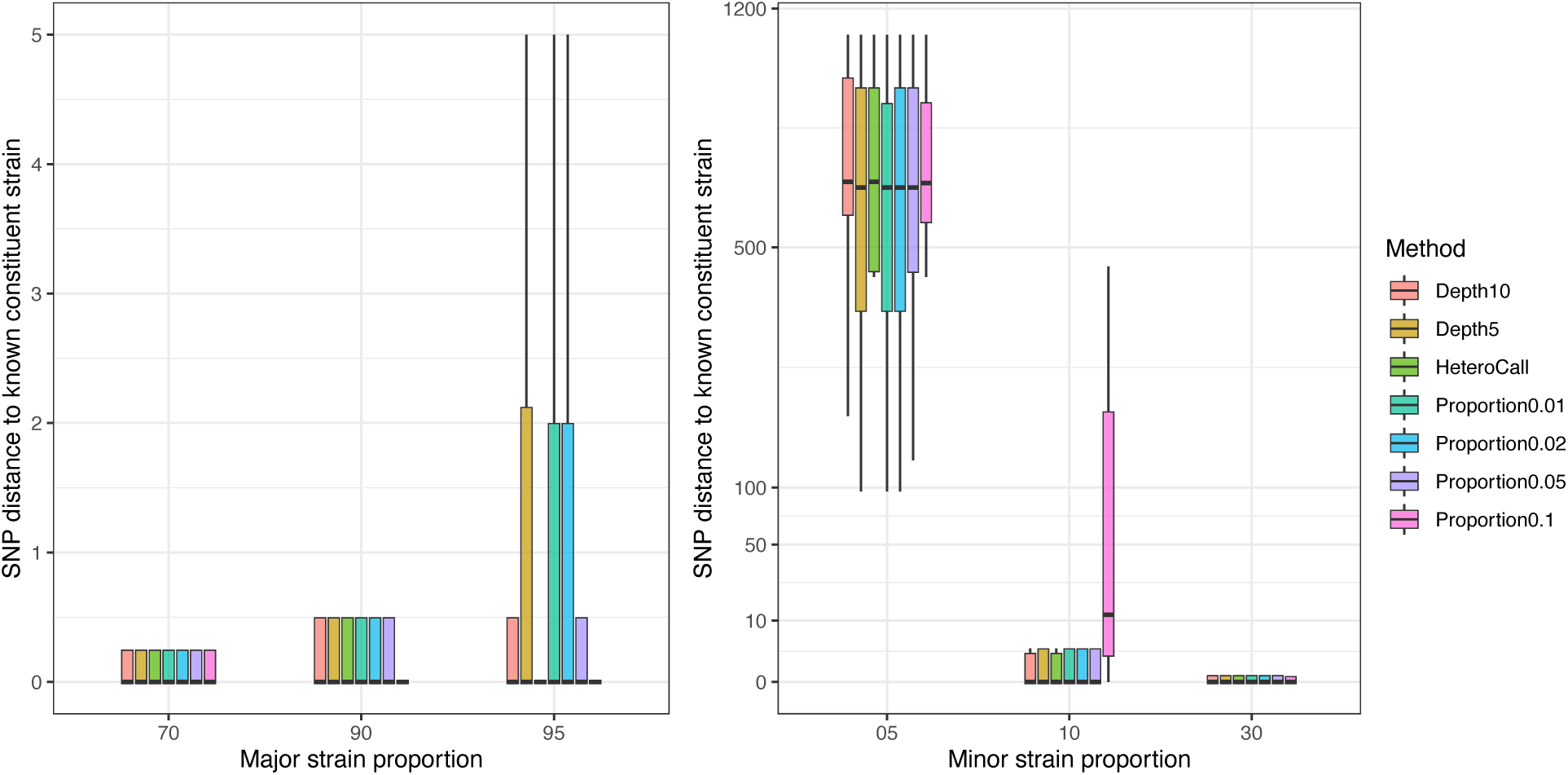
The distance between the closest identified strain and the known constituent strain using seven different approaches to characterizing hSNPs for the major strain sequence (A) and minor strain sequence (B) of the *in vitro* mixed samples.

**Supplementary figure S4.**
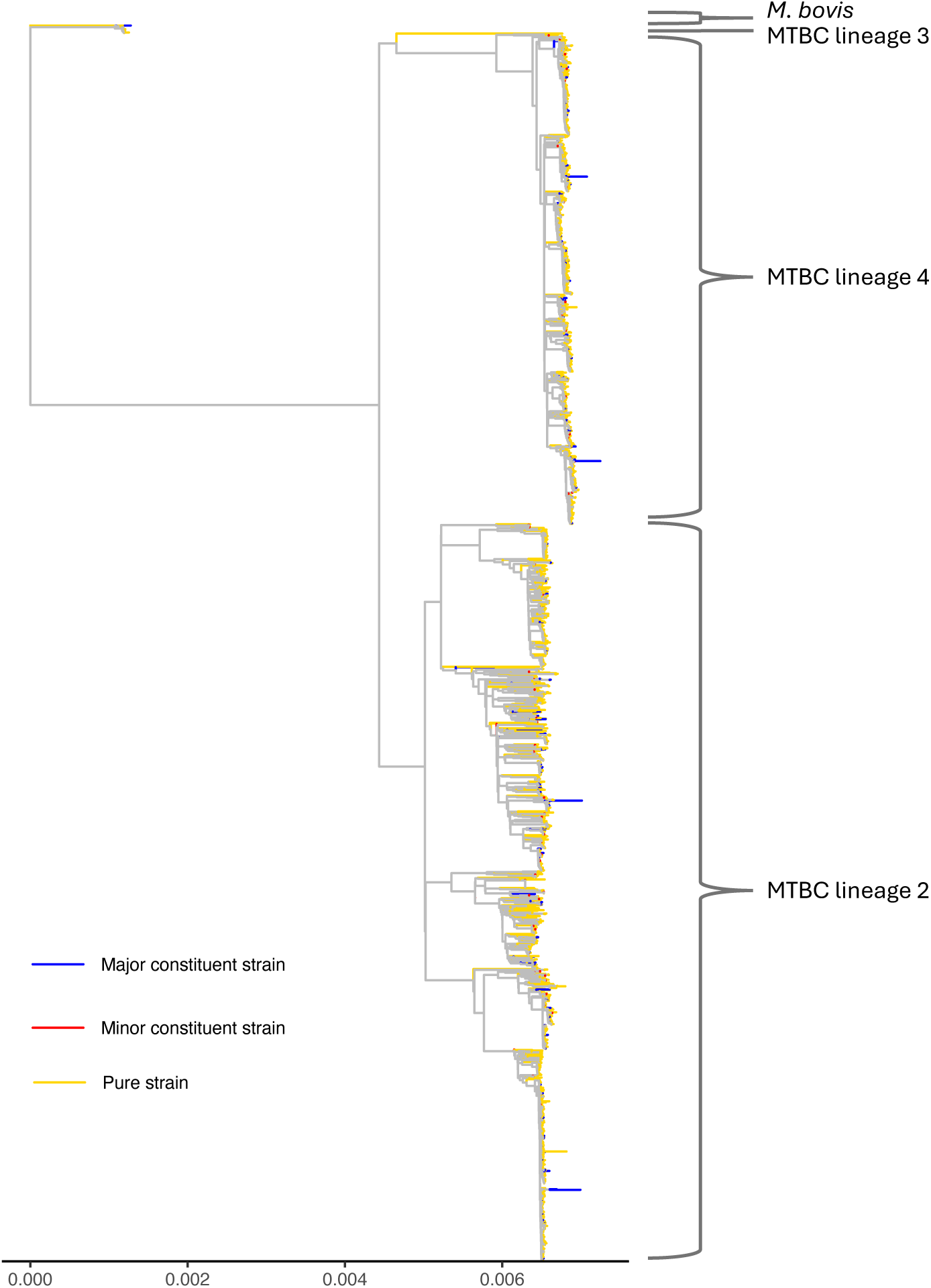
A maximum likelihood phylogenetic tree of the real-world Mtb samples from Moldova including the reconstructed constituent strains of predicted mixed infections. Terminal branches colored by the infection status of the sample at the tip, the non-mixed ‘pure’ strains in yellow, the major constituent strains of mixed infections in blue, and the minor constituent strains of mixed infections in red.

**Supplementary figure S5.**
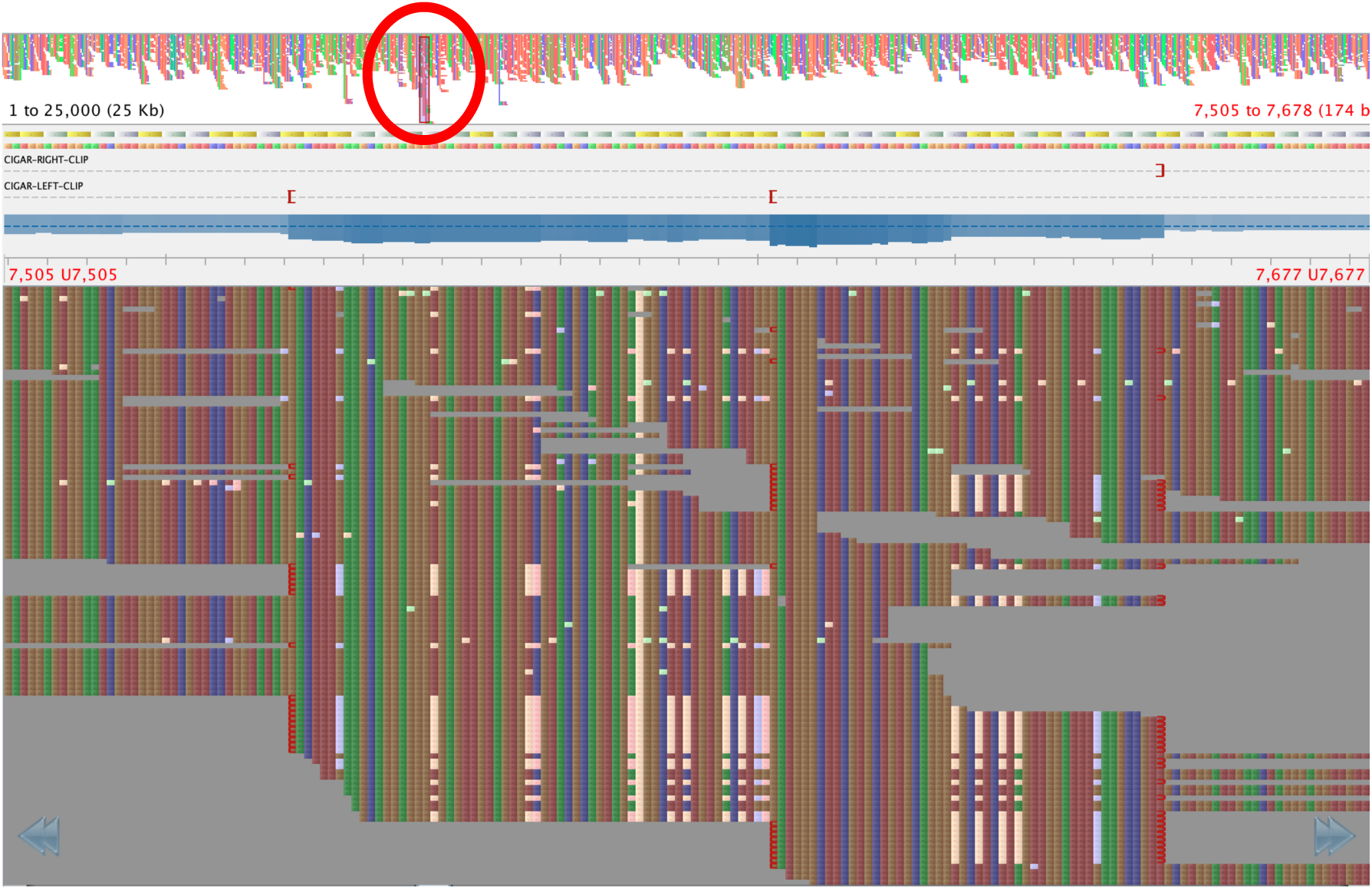
An example section of a binary alignment (BAM) file, visualized using Tablet ^31^, from a Moldova *Mtb* isolates (TB-222-6565-19_S218) that was not classified as mixed but harbored a high number of hSNPs. Variants found in aligned reads against the H37Rv reference sequences are highlighted, showing multiple hSNPs close in the genome and coverage almost double the average for surrounding regions, shown within the red circle.

